# Virulence constrains transmission even in the absence of a genetic trade-off

**DOI:** 10.1101/2021.10.07.463510

**Authors:** Diogo P. Godinho, Leonor R. Rodrigues, Sophie Lefevre, Laurane Delteil, André F. Mira, Inês R. Fragata, Sara Magalhães, Alison B. Duncan

## Abstract

The virulence-transmission trade-off predicts that parasite fitness peaks at intermediate virulence. However, whether this relationship is driven by the environment or genetically determined and if it depends on transmission opportunities remains unclear. We tackled these issues using inbred lines of the macro-parasitic spider-mite *Tetranychus urticae*. When transmission was not possible during the infection period, we observed a hump-shaped relationship between virulence and parasite fitness, as predicted by theory. This was environmentally driven, as no genetic correlation between traits was detected. However, when transmission to uninfected hosts occurred during the infection period, virulence was positively, environmentally and genetically correlated with parasite fitness. Therefore, the virulence-transmission trade-off depends on within-host dynamics and on the timing of transmission, rather than on a genetic correlation. This fundamental correlation may thus be easier to manipulate than previously thought.

## Introduction

Virulence, the harm inflicted by parasites on their hosts, is a trait with high relevance for human, animal, plant and ecosystem health. It is also an evolutionary puzzle, as by harming hosts, parasites seemingly jeopardize their chances of being represented in subsequent generations, that is, their fitness (Alizon et al., 2009).

The most widely accepted explanation for the existence of virulent parasites is the virulence-transmission trade-off hypothesis (Alizon et al., 2009; Anderson and May, 1982). According to this hypothesis, within-host growth is a component of parasite fitness (its reproductive growth rate, R_0_), but this is expected to entail high levels of virulence. High virulence, in turn, may lead to premature host death, hampering transmission, thus ultimately the growth of parasite populations (Anderson and May, 1982, 1979). Therefore, parasite fitness is expected to be maximised at intermediate virulence levels (Anderson and May, 1982). Despite the centrality of this hypothesis to the understanding of host-parasite interactions, and evidence of genetic variance for parasite traits (Little et al., 2008; Louhi et al., 2013; Mackinnon and Read, 1999a), whether the trade-off is due to genetic correlations among traits or is environmentally driven remains to be addressed. Disentangling these alternative possibilities is key to identify the conditions under which parasite traits can evolve independently, which could be applied on strategies for the management of parasite virulence.

Most studies in support of the trade-off hypothesis have used parasite isolates that differ genetically, but also in their recent ecological and evolutionary history, as they have different geographic origins (de Roode et al., 2008; Doumayrou et al., 2013; Ebert, 1994; Mackinnon and Read, 2003; Mackinnon and Read, 1999b). This may lead to spurious correlations between traits. Using inbred lines derived from the same parasite population allows this issue to be overcome. Additionally, most studies address the trade-off hypothesis by measuring transmission (or a proxy thereof) at the end of the infection period (Acevedo et al., 2019; Mackinnon and Read, 2003; Mackinnon and Read, 1999b), which mimics a parasite with a single transmission event. However, several parasites transmit continuously during the infection period (e.g. HIV, malaria (Fraser et al., 2007; Mackinnon and Read, 1999b)), and this may affect the relationship between virulence and transmission (Day et al., 2011). Therefore, addressing the generality of the virulence-transmission trade-off requires accounting for these different parasite life cycles.

Here, we tested whether the virulence-transmission trade-off was determined by genetic correlations and/or was environmentally driven using 15 inbred lines derived from one outbred population of the spider mite *Tetranychus urticae*, a plant macro-parasite (Godinho et al., 2020). Spider mites spend their entire life-cycle on their host plants (Helle and Sabelis, 1985) causing damage that correlates negatively with plant fitness (Fineblum and Rausher, 1995). This damage is visible and quantifiable through chlorotic lesions on the leaf surface ((Mira et al., 2021); Figure S1), which represents a reliable measure of virulence. Once mites become adult and mate, females either remain on the plant or they disperse and infect new hosts (Bitume et al., 2013). Spider mites have ambulatory and passive aerial dispersal, hence transmission can depend on environmental factors such as wind (Smitley and Kennedy, 1985). They can transmit during the infection period or overexploit the host plant before transmission occurs (Smitley and Kennedy, 1985). Transmission may, thus, be dependent on many factors such as within-host parasite density and/or the availability of suitable hosts to infect (Bitume et al., 2013; De Roissart et al., 2015), making this system ideal to test if such factors affect the virulence-transmission trade-off. Because mites are macro-parasites, we used the number of adult daughters produced in a host as a measure of parasite fitness, R_0_ (Anderson et al., 1986; May and Anderson, 1979). We assessed whether a relationship between virulence and R_0_ was determined by genetic differences among lines, and/or by the build-up of density-dependence within the host, by varying initial densities of infesting mites (infection dose). Additionally, we evaluated whether this relationship was affected by opportunities for transmission during the infection period in two separate experiments. In the first, all parasite life-history traits were measured on a single host patch and no transmission was allowed until the end of the infection period. In the second, the adult female offspring of the parasite could disperse continuously during the infection period to a new host patch.

## Results

### Genetic variation for virulence, R_0_ and transmission

If the virulence-transmission trade-off is to be driven by genetic correlations among parasite traits, genetic variance for these traits must be present in a parasite population. We thus measured the variance for virulence, parasite fitness and transmission among the *T. urticae* inbred lines, using per capita measurements, which allows broad sense heritability, *H*^*2*^, to be determined. We found significant genetic variance for all traits (Fig. S2). Additionally, all traits were affected by the initial density of founding females on the host patch. The exception was for per capita transmission to new host patches (in the experiment with continuous transmission) (Table S1). Broad sense heritability was significant for all traits measured, with levels similar among experiments (Table S2).

### Genetic and environmental correlations between parasite traits

#### Virulence and R_0_, transmission at the end of the infection period

We first assessed virulence and R_0_ in the absence of uninfected hosts to which parasites could transmit (i.e., mimicking a parasite life cycle with transmission at the end of the infection period only; Fig. S3): we infected hosts (bean leaf patches) with 5, 10 or 20 females from each inbred line for 4 days, then measured damage (virulence) and the number of females produced 10 days later (R_0_). In support of the trade-off hypothesis, we found a hump-shaped environmental relationship (resulting from the residual co-variance of the model only) between virulence and R_0_. This was shown by the model with a squared term between these traits having a lower DIC, as compared to the model with only the linear term (DIC = 3379 and 3382, respectively). The model with the lowest DIC included density, hence this factor affected trait correlations. This was corroborated by the analysis of trait correlations for each density separately, as we found a positive correlation at low density, no relationship at intermediate density, and a negative correlation at high density (Fig. 1a, Table S3). This suggests that the relationship between these traits is modulated by density dependence: beyond a certain level of virulence, within-host competition prevents more daughters from becoming adult, such that R_0_ is maximised at intermediate levels of virulence, as predicted by the trade-off hypothesis. We observed that females at higher densities do not lay fewer eggs (results not shown), suggesting that density-dependence operates during juvenile development. Evidence for density-dependence has been found in this (De Roissart et al., 2015; Rotem and Agrawal, 2003), and other host-parasite systems (Ebert et al., 2000; Pollitt et al., 2013).

**Figure 1.**
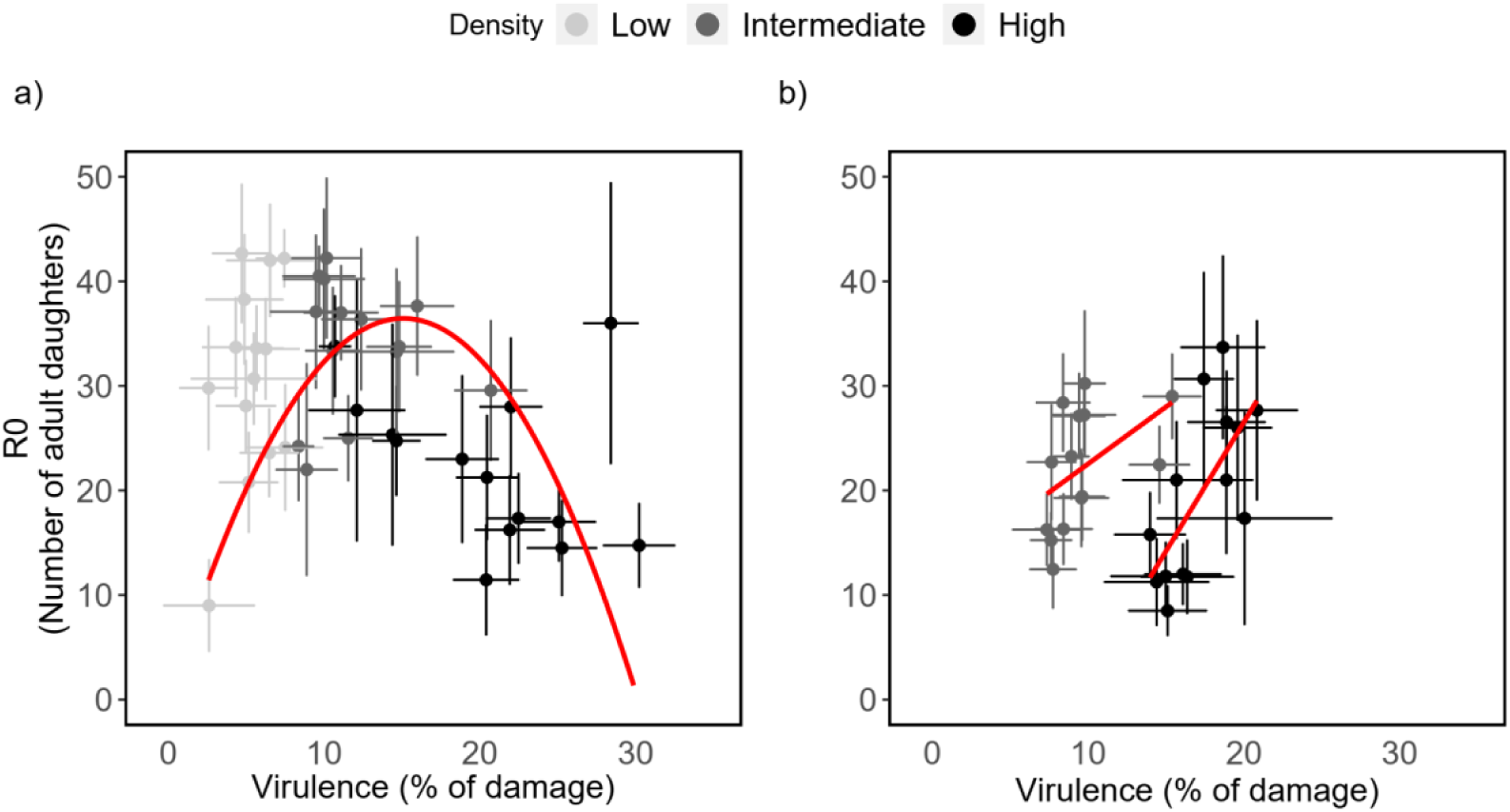
Correlation between virulence and the production of adult daughters in inbred lines of *T. urticae* infecting a host patch. The damage inflicted on the host patch (i.e., virulence) and the number of daughters produced (the parasite reproductive rate R_0_) were measured in a set-up with a) no uninfected hosts available; b) uninfected hosts available during the infection period. Shades of grey represent different densities; dots are the mean for each inbred line ± standard error; regressions are represented in red.

#### Virulence, R_0_ and transmission to uninfected hosts, continuous transmission

Next, for the two highest densities (10 and 20 females), we tested whether the presence of uninfected hosts (i.e. mimicking a parasite life cycle with continuous transmission during the infection period, Fig. S3) would modify the relationship between virulence and R_0_. Despite no overall environmental correlation between virulence and R_0_, this correlation was positive at both intermediate and high densities (Table S3; Fig. 1b). This suggests that the negative effects of high densities on R_0_ were alleviated by the possibility of moving to other hosts. Additionally, this correlation was affected by the inbred line identity, indicating that it has a genetic basis. Probably, in these conditions, more virulent genotypes suffer less from rapid host exploitation, because they can escape to new hosts. Moreover, we found a positive correlation between R_0_ and transmission to uninfected hosts (Table S3; Fig. 2), which is in accordance with theory (Anderson and May, 1982).

**Figure 2.**
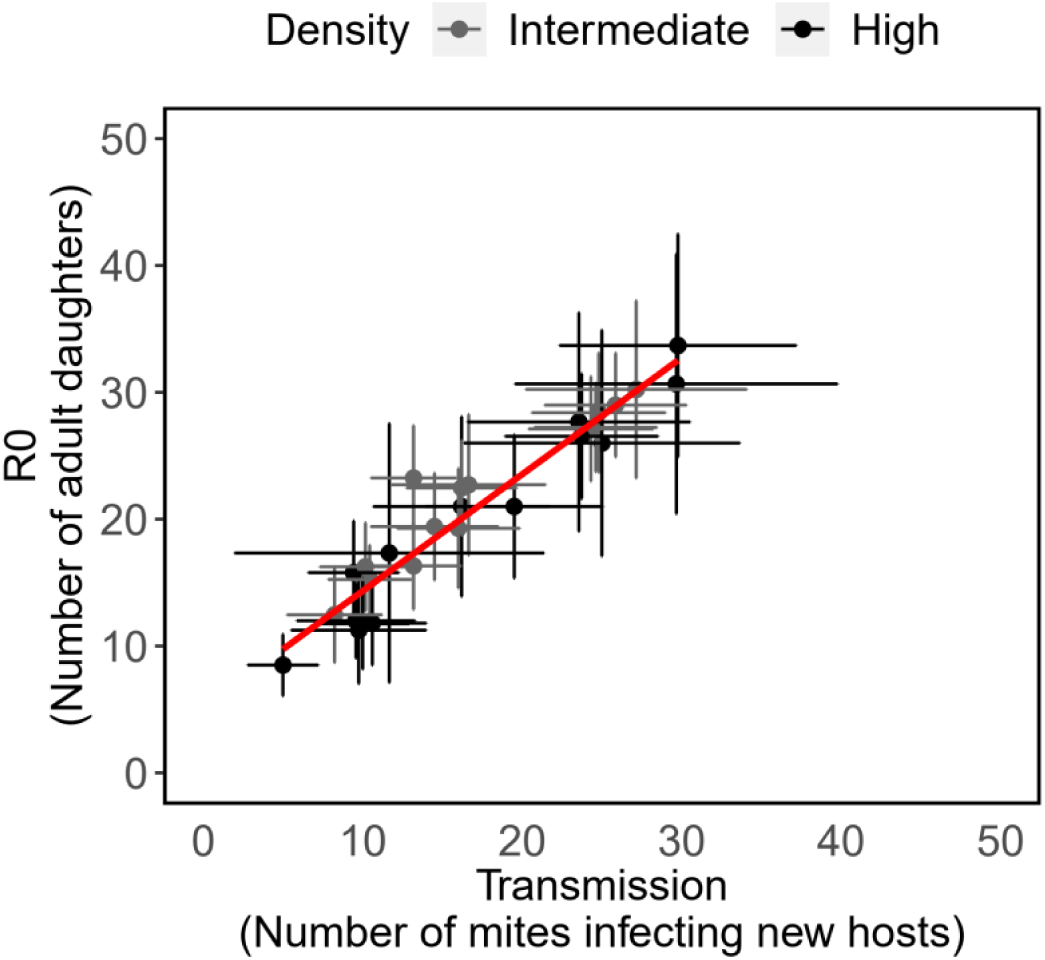
Correlation between the production of adult daughters and transmission in inbred lines of *T. urticae* infecting a host patch. The number of daughters produced (the parasite reproductive rate, R_0_) and transmission (the number of mites infecting new hosts) for inbred lines of *T. urticae* at different starting densities (Intermediate = 10 females; High = 20 females). Shades of grey represent different densities; dots are the mean for each inbred line ± standard error; the regression is represented in red.

## Discussion

In this study we show a hump-shaped relationship between virulence and transmission that does not have a genetic basis. Indeed, R_0_ is maximized at intermediate levels of virulence, as postulated in theoretical models (Anderson and May, 1982, 1979), but this is due to within-host density dependence. Moreover, this hump-shaped relationship disappears when transmission is continuous during the infection period. This may explain the mixed evidence for the occurrence of a trade-off in earlier studies and reinforces the idea of including the whole parasite life-cycle in the experimental set-up (Acevedo et al., 2019; Alizon and Michalakis, 2015). Indeed, if transmission timing affects the relationship between virulence and R_0_, failing to include this important step of the parasite life cycle in the experimental set-up may lead to conclusions based on incomplete evidence. Contrasting relationships between virulence and R_0_ reliant on transmission timings may have important consequences for ecology and evolution for parasites of the same or different species.

In parasites with transmission at the end of the infection period, the lack of a genetic correlation between virulence and R_0_ means that selection in one trait may not affect the other, potentially maintaining variance for both traits, as observed in this (Fig. S2) and other systems (Dutta et al., 2021; Little et al., 2008; Louhi et al., 2013; Mackinnon and Read, 1999a). This may enhance the ability to cope with variability in host populations (Dutta et al., 2021; Nørgaard et al., 2021), which is particularly relevant for generalist parasites, such as *T. urticae*. In the absence of a genetic link with virulence, transmission may instead vary with other factors such as host availability and variability (King and Lively, 2012; Parsche and Lattorff, 2018). Conversely, selection on virulence may depend on other epidemiologically related traits, such as co-infections or the host immune system (Alizon et al., 2009). In parasites with continuous transmission, the positive genetic correlation between virulence and R_0_ suggests there should be selection for higher virulence. If this is the case, why then do we still find genetic variance for this trait? We propose two non-mutually exclusive hypotheses. First, *T. urticae* is a generalist parasite, hence optimal virulence may vary with the host species (Rioja et al., 2017). Second, transmission within the infection period relies on the occurrence of hosts to which parasites can transmit. This is not necessarily always possible, as uninfected hosts may be locally absent or they may become rapidly infested (Crossan et al., 2007; Hochberg, 1991). Although mites may be passive dispersers, they base their decision to leave a host patch on their perception of cues (volatiles) from hosts in the environment, including their infection status (Kiedrowicz et al., 2017; Pallini et al., 1997). Therefore, if there are no uninfected hosts mites may remain on their host, which will eventually result in them switching to a parasite life cycle without continuous transmission. Thus, variability in transmission timing, may contribute to the maintenance of genetic variance for virulence.

The difference in parasite life cycles analyzed here has obvious implications for the virulence-transmission trade-off that have not been shown empirically before. Still, our experimental design did not include variation in the availability and/or heterogeneity of recipient hosts. Considering these host population characteristics would further contribute to our knowledge about how differences in parasite life cycles may affect the virulence-transmission trade-off and, therefore, influence disease dynamics.

## Materials and Methods

### Biological system

*Tetranychus urticae* is an ectoparasite of over 1000 host plant species (Rioja et al., 2017). Females lay eggs on the leaf surface that hatch up to 4 days later. There are three immature stages, punctuated by quiescent stages, after which the mites become adult (∼9 days from hatching to adulthood in our laboratory), with their complete life cycle occurring on their host plant (Helle and Sabelis, 1985). They feed by injecting their stylet into plant cells, mostly parenchyma cells, and sucking out the cytoplasm, producing chlorotic lesions in the form of white spots ((Mira et al., 2021), Figure S1). The damage inflicted by *T. urticae* (our measure of virulence), together with high intrinsic growth rates, has important consequences for plant growth and yield, resulting in major economic losses worldwide (Helle and Sabelis, 1985).

### Populations used

*Tetranychus urticae* were collected on different host plants, in Portugal in 2013 (Zélé et al., 2018), and have since been reared on bean plants (*Phaseolus vulgaris*, variety Prelude), at the University of Lisbon. In October 2015, 50 individuals from 6 different field populations (total: 300) were collected and mixed to form an outbred population that has been kept at high densities (>1000). In October 2016, inbred lines were created from this population by sib mating. This procedure was repeated for 14 generations, ensuring an inbreeding coefficient above 94% (Godinho et al., 2020). Inbred lines allow simultaneous measurement of many individuals of the same (nearly) homozygous genotype, thus increasing the accuracy of genetic estimates (Godinho et al., 2020). Because they were derived from the same population, all lines share the same evolutionary and environmental history. Additionally, given that this population was outbred, genetic variation across lines is expected to be high (Godinho et al., 2020). Lines were maintained separately on bean leaf patches in Petri dishes. A subset of 15 inbred lines were transferred to the University of Montpellier in January 2018 and maintained on bean leaves (variety Pongo) in small plastic boxes (255 mm length x 183 mm width x 77 mm height) at optimal conditions (25°C with a 16:8 L: D cycle, at 60% relative humidity). These same conditions were kept throughout all experiments.

Prior to the experiments, cohorts of spider mites from each inbred line were created by isolating 40 to 50 mated females of each line on bean leaves placed on water-saturated cotton wool in boxes. These females laid eggs for 48h. Fourteen days later, mated females (daughters) were used in the experiments. Not all inbred lines are represented in each experiment due to too few individuals available at the start of the experiment (N between 12 to 14 lines).

### Virulence and R_0_, transmission at the end of the infection period

Females of each inbred line were randomly assigned to a low, intermediate or high-density treatment, corresponding to 5, 10 or 20 founding females, respectively, on a 4 cm^2^ bean leaf patch, placed on wet cotton wool in plastic boxes. All females were allowed to feed and lay eggs on their leaf patches for 4 days. After this period, adult females were killed, and a photograph of each patch was taken using a Canon EOS 70D camera. The amount of damage inflicted by spider mites was measured using ImageJ (Schneider et al., 2012) and Ilastik 1.3 (Sommer et al., 2011). Briefly, the background from each photo was removed in ImageJ, then we distinguished damaged area from healthy leaf using Ilastik and finally the damaged area was calculated using the colour contrast between damaged and undamaged leaf tissue in ImageJ (Fig. S1, (Mira et al., 2021)). Control leaf patches, never exposed to spider mites, were placed in the experimental boxes for the same time period and photographed. These were used to establish an average baseline “damage”, which was subtracted from each measurement, to provide an estimate of virulence. After a period of 14 days, the adult daughters surviving on each patch were counted. In this set-up, transmission would only be possible after this measurement, i.e., at the end of the infection period. There were 3 to 13 replicates for each inbred line per density, distributed across 3 blocks.

### Virulence, R_0_ and transmission to uninfected hosts, continuous transmission

Adult females were randomly assigned to the intermediate or the high-density treatments (10 or 20 females, respectively, on a 4 cm^2^ bean leaf patch placed on water saturated cotton wool). As in the previous experiment, females were left to lay eggs for 4 days, after which they were killed, and a photograph was taken of each leaf to measure the damage inflicted (Fig. S1). On day 4, a second leaf patch, uninfected by spider-mites, was placed beside the first and connected to it by a 3 × 1 cm Parafilm bridge (Fig. S3). In this way, the emerging adult female offspring could walk across this bridge and infect a new leaf patch. The number of adult daughters on the new host patches was checked on days 11, 12 and 13 (in block 1 only on days 12 and 13). When there were more than 15 offspring on the new patch, the latter was replaced by a new one. Host patches were replaced so that uninfected patches were always available. On day 14, we counted the number of adult daughters on the original host patch and on each of the new patches. Transmission was inferred by the cumulative number of females that infected a new host patch. This set-up mimics the life cycle of a parasite with continuous transmission during the infection period, as found in several systems (Fraser et al., 2007; Mackinnon and Read, 1999b). There were 5 to 16 replicates for each inbred line per density treatment, distributed across 4 blocks.

### Statistical analysis

We present correlations between *T. urticae* life-history traits: damage inflicted (a measure of virulence), adult daughters produced (a measure of R_0_ for macro-parasites (Anderson et al., 1986; Anderson and May, 1979)) and the number of females infecting a new host (a measure of transmission). We consider total values per host patch (Table S3), which are generally used in theoretical models (Anderson et al., 1986; Anderson and May, 1982; Day et al., 2011; May and Anderson, 1979) and correspond to the traits measured in most experimental studies testing virulence-transmission correlations (de Roode et al., 2008; Doumayrou et al., 2013; Mackinnon and Read, 2003, 1999b). Genetic and environmental correlations between virulence and R_0_, and between R_0_ and transmission in the experimental set-up mimicking continuous transmission, were performed using a multi-response generalized linear mixed model fitted with an MCMCglmm (package MCMCglmm (Hadfield, 2010)). Genetic correlations were determined by including the identity of the line as a random factor in each model and assessing the highest posterior density interval (HPDI) of the genetic (G) structure of the model, which represents the (co)variances between the two traits evaluated across inbred lines (Hadfield, 2010). Environmental correlations were obtained by assessing the HPDI of the residual (R) structure in the same model (Hadfield, 2010). Effects were considered significant when the HPDI did not overlap with 0. The effect of density on each correlation was assessed by comparing the deviance information criterion (DIC) of the models including density as a fixed factor or not. We also report the genetic and environmental correlations when considering each density level separately (Table S3). In addition, we tested whether a non-linear regression might best describe the environmental relationship between virulence and R_0_ in the experiment mimicking transmission at the end of the infection period only. To this aim, we compared two MCMCglmm models both with R_0_ fitted as the response variable and inbred line and block as random factors. Both models included density as a fixed factor and virulence as a covariate, with one model including only the linear term for virulence and the other both the linear and the quadratic terms.

For the assessment of genetic variance (variance among inbred lines) and the effect of density on this variance we used per capita values, by dividing the value for each host patch by the initial density of females, as these values are more representative of individual variation (Fig. S2, Table S2). We then applied generalized linear mixed models fitted with a Markov Monte Carlo Chain approach (Hadfield, 2010). Broad-sense heritability, 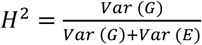 (Falconer and Mackay, 1996) and the corresponding confidence intervals were extracted from the abovementioned models for each trait.

All models initially included 300000 iterations, with a burn-in of 10000 iterations, thinning of 100 and a flat prior: For GLMMs (to assess the genetic variance within a trait), V= 1 and nu=0.002; For Multi-response GLMMs (to assess trait correlations), V= matrix(c(1,0,0,1), ncol= 2, nrow= 2) and nu= 0.002. Flat priors were used to allow the hyper-parameter values to reflect a reasonable range of values for the traits in question, without any previous information about them or their co-variance. All models were checked for convergence with a stationary test using the heidel.diag function and for autocorrelations for the Markov chain within fixed and random terms using the autocorr.diag function. When models failed one of these tests, the number of iterations was increased to 500000 or to 700000 and the burn-in to 20000 or 50000. All figures were produced with the ggplot2 package in R and the regressions included were fitted with the geom_smooth function (Wickham and Winston, 2016).

## Acknowledgments

We deeply thank Élio Sucena, Flore Zélé, Margarida Matos, Oliver Kaltz, Sylvain Gandon and Yannis Michalakis, for their very useful suggestions and comments.

## Author contributions

Conceptualization and supervision by SM and ABD; Investigation by DPG, ABD, LRR, SL, AFM and LD; Formal analysis by DPG and IRF; Visualization by DG, LRR and IRF; Writing the original draft by DPG, SM and ABD with reviewing and editing by LRR and IRF.

## Data and materials availability

Data will be deposited in Figshare upon acceptance.

## Supplementary Information for

**Figure S1.**
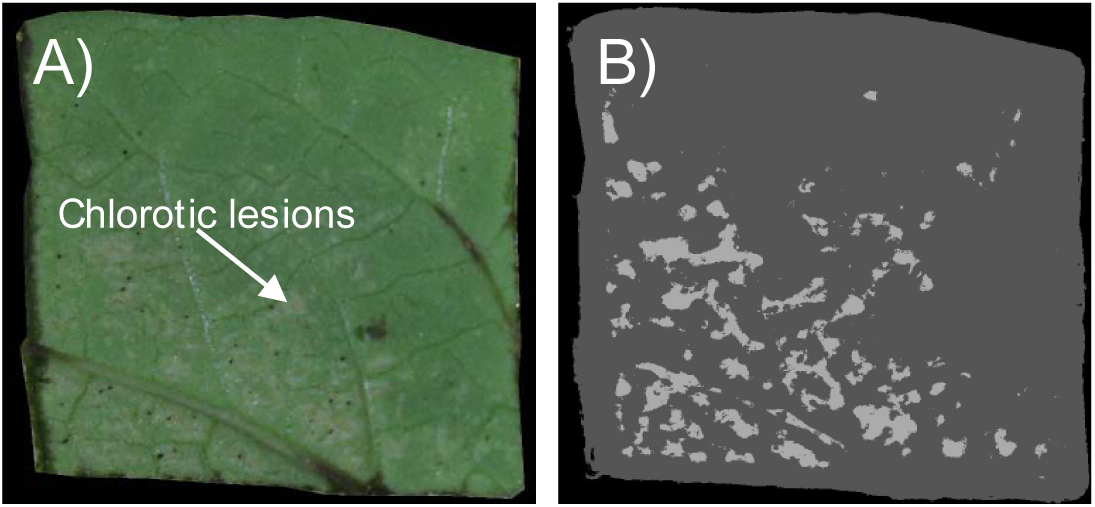
Example of a host patch cut from a bean plant (*Phaseolus vulgaris*) upon image acquisition and after software output. A) Leaf damage (chlorotic lesions) caused by *T. urticae* feeding. B) The photograph is transformed into a simple segmentation image using Ilastik 1.3. Areas of leaf are damage shown in light grey and correspond to our measure of virulence.

**Figure S2.**
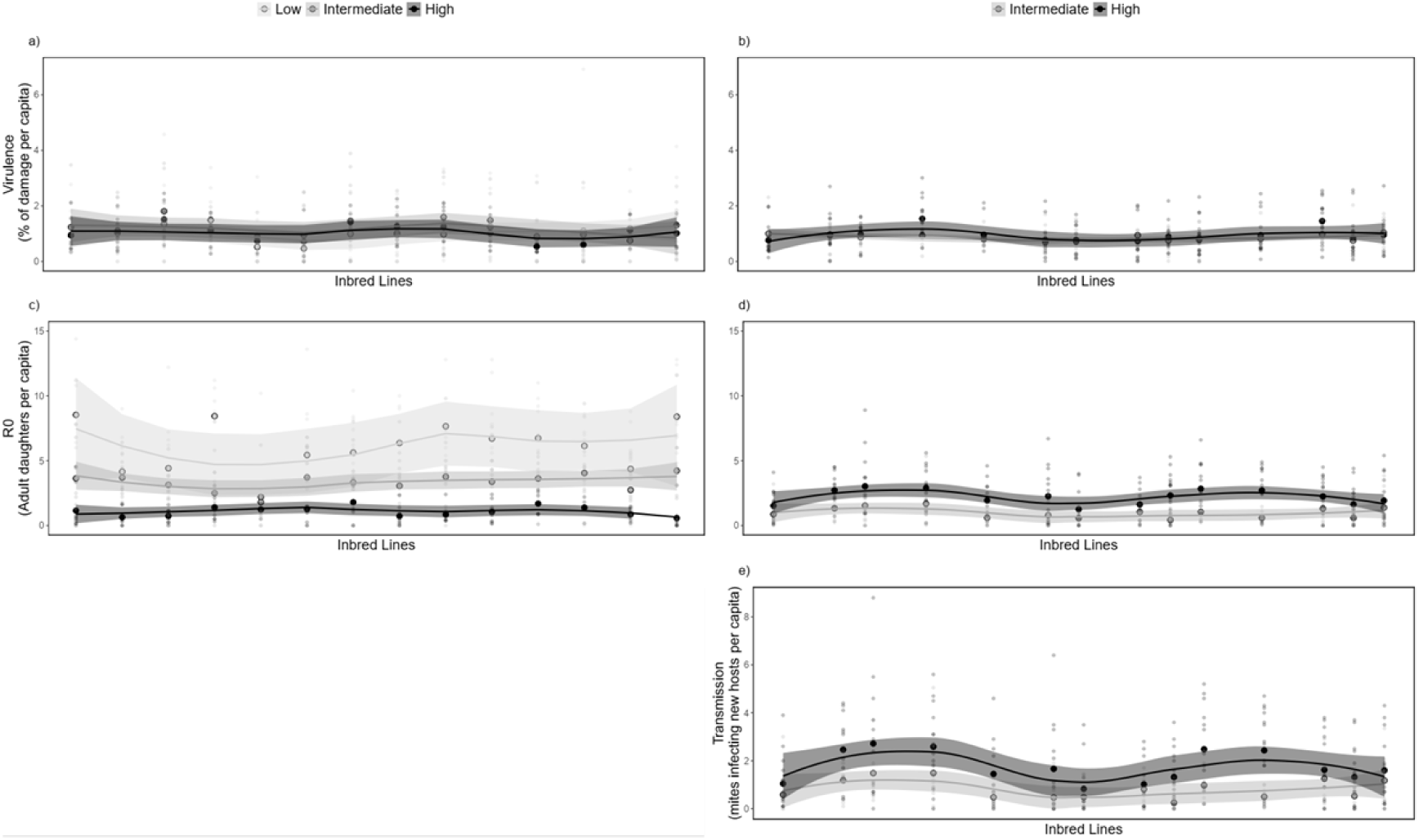
Genetic variance. Genetic variance for virulence (% of damage inflicted; a and b), R_0_ (the number of adult daughters; c and d), and transmission (number of mites infecting new hosts; e), measured per capita in the different experiments (Experiment 1 – a and c; Experiment 2 – b, d, and e). Large circles represent the average per line, per density (light grey = Low density; dark grey = Intermediate density; black = High density). The shape of the curve for each density (light grey = Low density; dark grey = Intermediate density; black = High density) was calculated using a polynomial regression fitted with the geom_smooth function.

**Figure S3.**
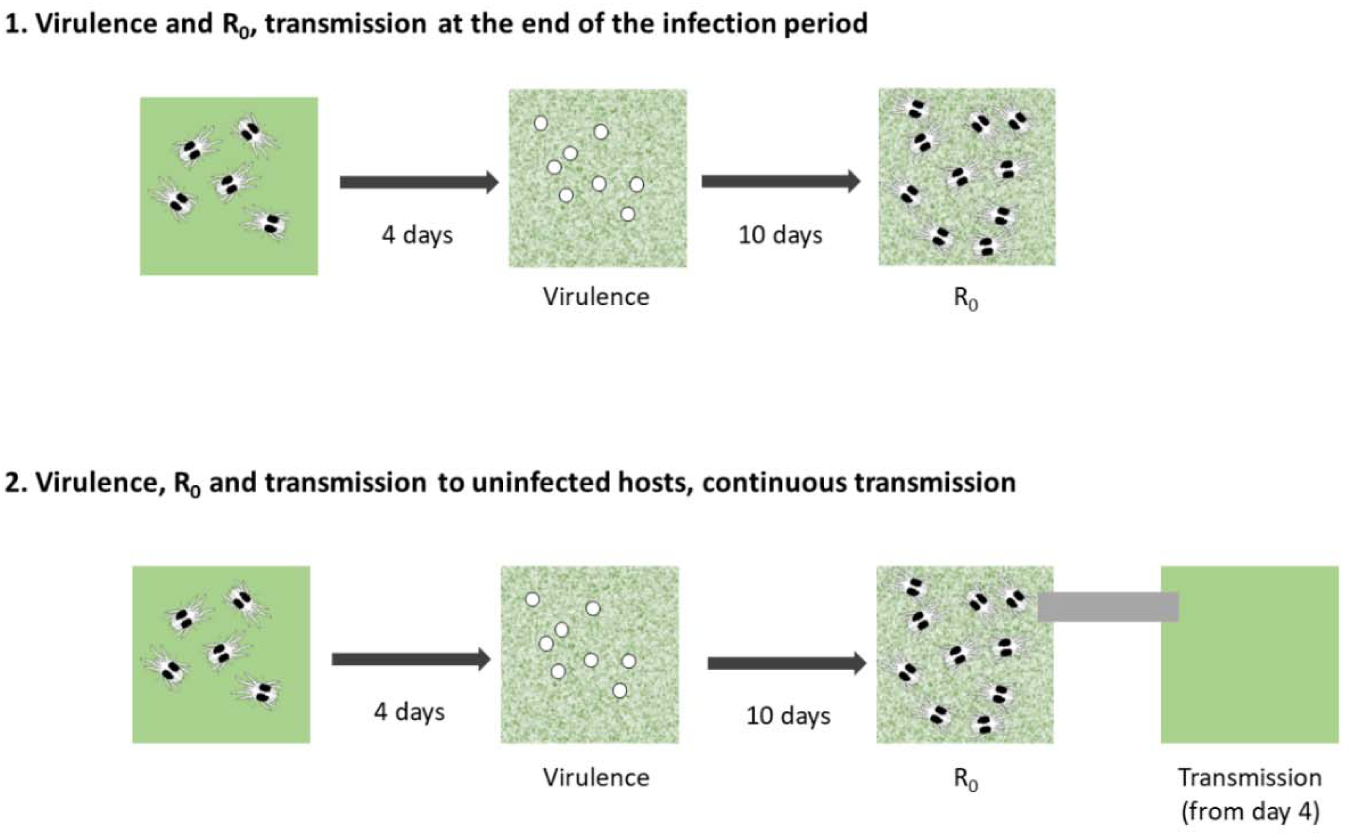
Experimental set-ups. Schematic representation of the experimental set-ups and the traits that were measured in each experiment. Day 0 shows 5 spider mites on a healthy 4cm^2^ leaf patch (low density treatment). On day 4, virulence (mottled white areas on leaf patches) was measured in experiments 1 and 2. R_0_ (i.e. number of adult daughters) was measured 14 days after mite installation. In experiment 2 transmission to an uninfected leaf patch was possible from day 4 to day 14.

**Table S1.**
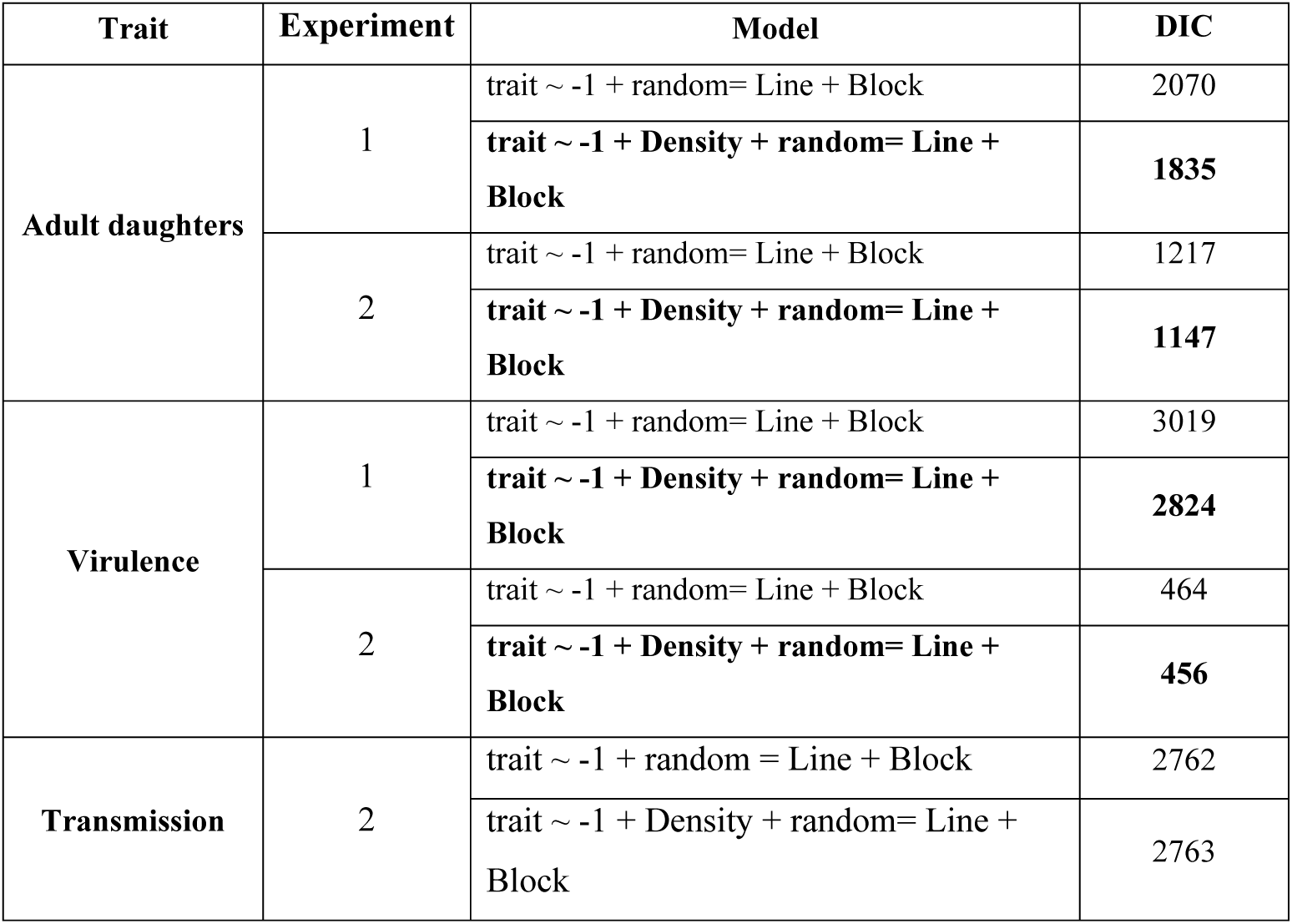
The effect of the initial density of female spider mites on the patch on the per capita production of adult daughters (R_0_), virulence and transmission (i.e. “trait” in the model column). Deviation information criterion (DIC) of the models (MCMCglmm package) measured in the different experiments with density included or not as a fixed factor. When applicable, best fit models are in bold. Experiment 1: transmission at the end of the infection period only; experiment 2: continuous transmission.

**Table S2.**
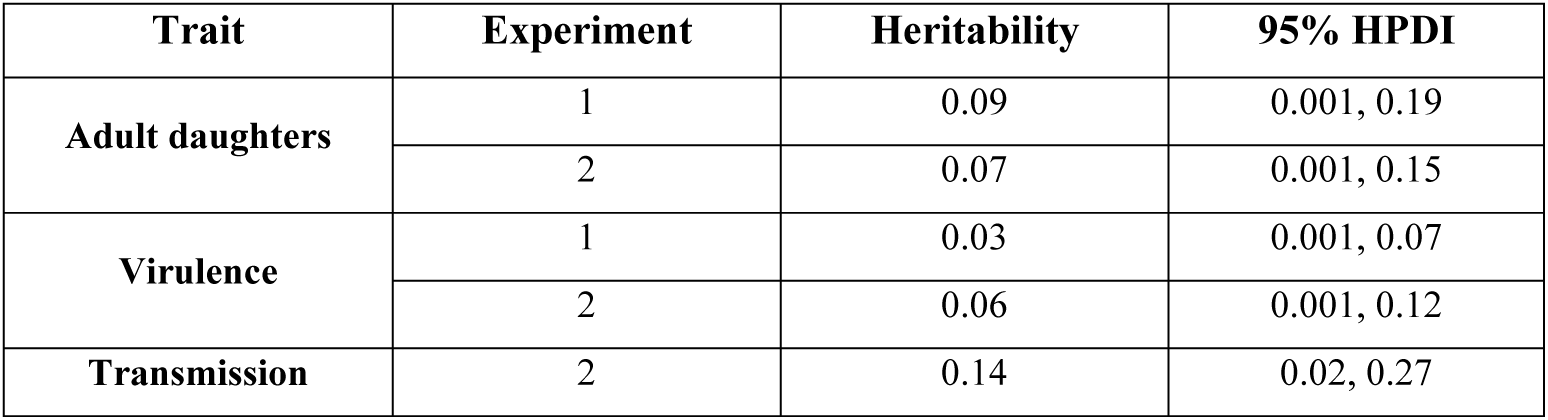
Trait heritability. Broad-sense heritability for the per capita production of adult daughters (R_0_), virulence and transmission measured in the different experiments. 95% highest posterior density intervals (HPDI) intervals for the heritability of each trait. All traits have significant heritability as no interval includes zero. Experiment 1: transmission at the end of the infection period only; experiment 2: continuous transmission.

| **Trait** | **Experiment** | **Heritability** | **95% HPDI** |
| --- | --- | --- | --- |
| **Adult daughters** | 1 | 0.09 | 0.001, 0.19 |
|  | 2 | 0.07 | 0.001, 0.15 |
| **Virulence** | 1 | 0.03 | 0.001, 0.07 |
|  | 2 | 0.06 | 0.001, 0.12 |
| **Transmission** | 2 | 0.14 | 0.02, 0.27 |

**Table S3.**
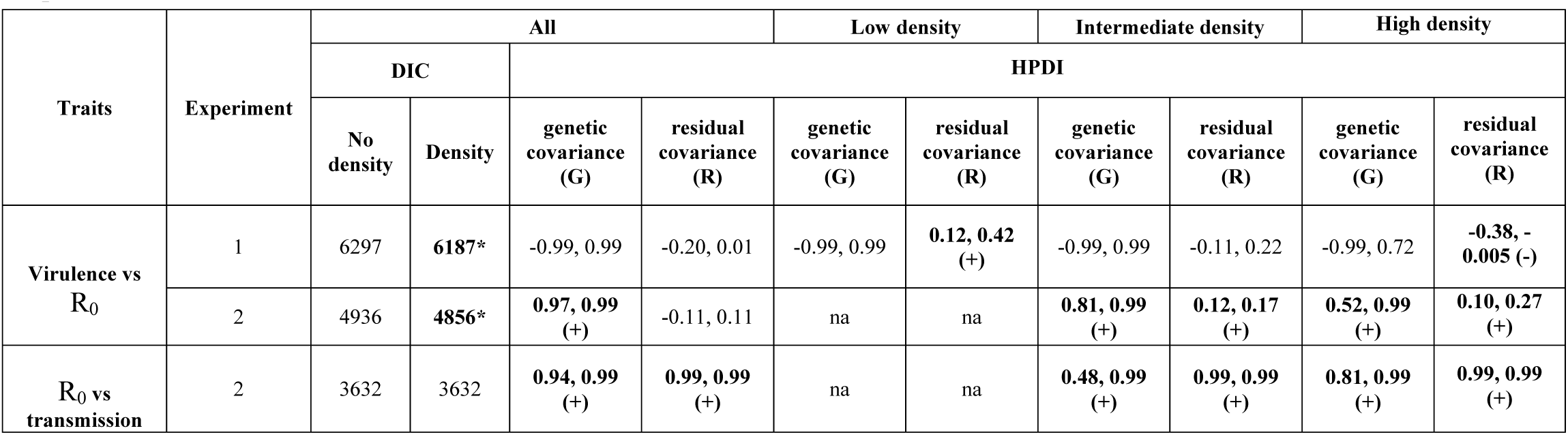
Genetic and environmental correlations. Genetic and environmental correlations - extracted from the genetic (random) or residual error structure of the models, respectively - between the production of adult daughters (R_0_) and virulence or transmission. All traits were measured per host (total value). The deviance information criterion (DIC) of models for all data with and without densities are shown. Highest posterior density intervals (HPDI) are shown for the model including all data (* model with lowest DIC, or in the case of R_0_ vs. transmission, the simplest model), and separately at the different densities. Intervals not overlapping 0 are shown in bold. The direction of the correlations is shown in brackets. Experiment 1: transmission at the end of the infection period only; experiment 2: continuous transmission.

| Traits | Experiment | All |  |  |  | Low density |  | Intermediate density |  | High density |  |
| --- | --- | --- | --- | --- | --- | --- | --- | --- | --- | --- | --- |
|  |  | DIC |  | HPDI |  |  |  |  |  |  |  |
|  |  | No density | Density | genetic covariance (G) | residual covariance (R) | genetic covariance (G) | residual covariance (R) | genetic covariance (G) | residual covariance (R) | genetic covariance (G) | residual covariance (R) |
| Virulence vs R<sub>0</sub> | 1 | 6297 | 6187* | -0.99, 0.99 | -0.20, 0.01 | -0.99, 0.99 | 0.12, 0.42 (+) | -0.99, 0.99 | -0.11, 0.22 | -0.99, 0.72 | -0.38, -0.005 (-) |
|  | 2 | 4936 | 4856* | 0.97, 0.99 (+) | -0.11, 0.11 | na | na | 0.81, 0.99 (+) | 0.12, 0.17 (+) | 0.52, 0.99 (+) | 0.10, 0.27 (+) |
| R<sub>0</sub> vs transmission | 2 | 3632 | 3632 | 0.94, 0.99 (+) | 0.99, 0.99 (+) | na | na | 0.48, 0.99 (+) | 0.99, 0.99 (+) | 0.81, 0.99 (+) | 0.99, 0.99 (+) |

